# Utilizing Pretrained Vision Transformers and Large Language Models for Epileptic Seizure Prediction

**DOI:** 10.1101/2024.11.03.621742

**Authors:** Paras Parani, Umair Mohammad, Fahad Saeed

**Affiliations:** Knight Foundation School of Computing and Information Sciences, Florida International University, Miami, FL, USA

**Keywords:** Large Language Model (LLM), Vision Transformer (ViT), Electroencephalography (EEG), Epilepsy, Seizure Prediction

## Abstract

Repeated unprovoked seizures is a major source of concern for patients suffering from epilepsy. Predicting seizures before they occur is of interest to both machine-learning scientists as well as clinicians, and is an active area of research. The variability of EEG sensors, type of seizures, and specialized knowledge required for annotating the data complicates the large-scale annotation process essential for supervised predictive models. To address these challenges, we propose the use of Vision Transformers (ViTs) and Large Language Models (LLMs) that were originally trained on publicly available image or text data. Our work leverages these pre-trained models by refining the input, embedding, and classification layers in a minimalistic fashion to predict seizures. Our results demonstrate that LLM’s outperforms the ViT’s in patient-independent seizure prediction achieving a sensitivity of 79.02% which is 8% higher compared to ViT’s and about 12% higher compared to a custom-designed ResNet-based model. Our work demonstrates the successful feasibility of pre-trained models for seizure prediction with its potential for improving the quality of life of people with epilepsy. Our code and related materials are available open-source at: https://github.com/pcdslab/UtilLLM_EPS/

## I. Introduction

Epilepsy is a chronic neurological disorder that causes unprovoked repeat seizures, and affects over 65 million people globally [1]. A seizure is a heightened activity of the brain that causes various symptoms ranging from simple loss of focus to loss of muscular control and unconsciousness resulting in falls and injuries [2]. Development of machine-learning models that utilize electroencephalography (EEG) data, and predict epileptic seizure is an active area of research. Successful ML models can enable patients and caregivers to avoid the significant financial and social costs related to seizures. Training generalizable models from scratch is challenging due to limited annotated data that might be available specific to different EEG sensors, epilepsy types, types of seizures, and co-morbid factors. The existing seizure prediction models, therefore, usually report results that are patient-specific on small datasets such as the CHB-MIT dataset [3]. Insufficient performance [4] with either low sensitivity or too many false positives are a well-known problem that requires new training and benchmarking approaches [5].

In this paper, we design and develop a strategy that can be used for fine-tuning models - for predicting seizures - that were originally trained on other data modalities. For this work, we have focused on investigating vision transformers (ViTs) and Large language models (LLMs). A typical epileptic seizure has 4 stages: *preictal* (before a seizure), *ictal* (main symptomatic phase), *post-ictal* (recovery stage) and *interictal* (periods of normal function between two seizures). Seizure prediction is a challenging task which involves discrimination of preictal and interictal segments, as compared to only detection (distinguishes ictal phases from the non-ictal phases). In this work, we solve two computational challenges: (1) how to refactor the data for models trained on images (ViT), and text (LLM) into a model that can process complex multi-channel EEG data which is then represented as time-series classification (TSC); (2) how to minimally retrain these models that result in better performance metrics as compared to models that are trained from scratch. To this end, the contributions in this paper are listed below:

- We propose wrangling methods to ***transform*** multi-dimensional time-series EEG data into a format suitable for ViTs (trained on images) and LLMs (trained on text).
- We ***re-design*** the final stage of the pretrained ViTs and LLMs by modifying (or adding in LLM) the classifier head to perform seizure prediction from this complex time-series data.
- We develop and implement a minimalist ***re-training*** strategy that requires unfreezing of a few pre-trained layers, while exhibiting superior results as compared to other state-of-the-art methods.
- We validated our strategy on two ViTs and one LLM on one of the ***largest patient-independent datasets*** for scalp EEG data [5].

The paper is organized as follows: Section II briefly discusses the related work. Section III describes the methods employed to make the pre-trained ViTs and LLMs ready for the seizure prediction task using EEG data, and concisely describes the computing resources and datasets to test the models. Section IV discusses the experimental results and provides recommendations for best practices and finally, Section V concludes the paper and provides future directions.

## II Related Work

We have identified the following four gaps in existing literature. In this study, we are designing and developing strategies and methods that can fill some of these gaps.

Gap 1. Non generalizability of existing models: One of the most successful end-to-end seizure prediction systems was designed in [6] that achieved a prediction sensitivity of 99.6% using a combined CNN-LSTM model with neural architecture search (NAS). Other recent works have used CNN-LSTM [7], 3D CNN-LSTM [8], and contrastive learning techniques [9] with a maximum sensitivity of 95%. However, the work of [4] demonstrated that the prediction sensitivity of multiple such models including [6] falls below 70% once leave one seizure out (LOSO) cross-validation (CV) - which prevents data leakage - is applied.

Gap 2. Transformers not used in an end-to-end configuration: To improve the performance of predictive models, a few recent works have applied the transformer architecture [10], [11]. Though originally only proposed for NLP tasks [12], transformers have been successfully adopted for other domains such as the vision transformer (ViT) [13]–[15]. Though transformers have been used as a second layer of feature extraction [11] [10] for seizure prediction, their usage in an end-to-end configuration is missing from the literature.

Gap 3. No patient-independent models exist or are tested on small cohorts: All of the above referenced works [6]–[11] test the models using a very small-sized cohort, and perform *only* patient-specific seizure prediction. More recent patient-independent models [16], [17] use a small cohort and apply CV that may result in data-leakage in training and validation. For effective assessment, datasets with disjoint training and testing sets as illustrated in our recent MLSPred-Bench benchmark [5] are needed.

Gap 4. Absence of Large language (LLMs) for seizure prediction: To the best of our knowledge, LLMs have not been exploited for solving the seizure prediction problem.

## III Materials and Methods

To assess the usability of pre-trained ViTs and LLMs for seizure prediction, we test two ViTs: ViT-1 is based on the Swin architecture [13], [14] and ViT-2 is based on the SegFormer [15]. One LLM based on the LongFormer [12] is also tested. Originally, ViT-1 and ViT-2 were used for image classification and semantic segmentation respectively. Typically, a ViT splits an image into fixed size patches, then applies linear embeddings to each patch and adds a positional encoding followed by a transformer block. Though multi-channel EEG time-series data is similar to an image in structure, it also has a temporal component along the x-axis in addition to the spatial component (channels) along the y-axis. Further, the fast sampling rates of EEG make the temporal axis more dominant which is very different to colored square-like images on which ViTs are trained.

For example, the EEG data used in this project is available as 12 ML-ready benchmarks [5] which contain 5-second segments labeled as either preictal or interictal. The EEG data comprises 20 channels sampled at 256 Hz which results in each segment being of shape 20 × 1280. Hence, an innovative strategy is needed to make the data ready for for ViTs. Our proposed overall strategy is shown in Fig 1. The rationale behind this strategy is to make multi-channel time-series data suitable for use with ViTs which have been designed to provide optimal performance on square-like 3-channel images. The need is justified because this approach best utilizes the billions of parameters pre-trained on terabytes of data.

**Fig. 1.**
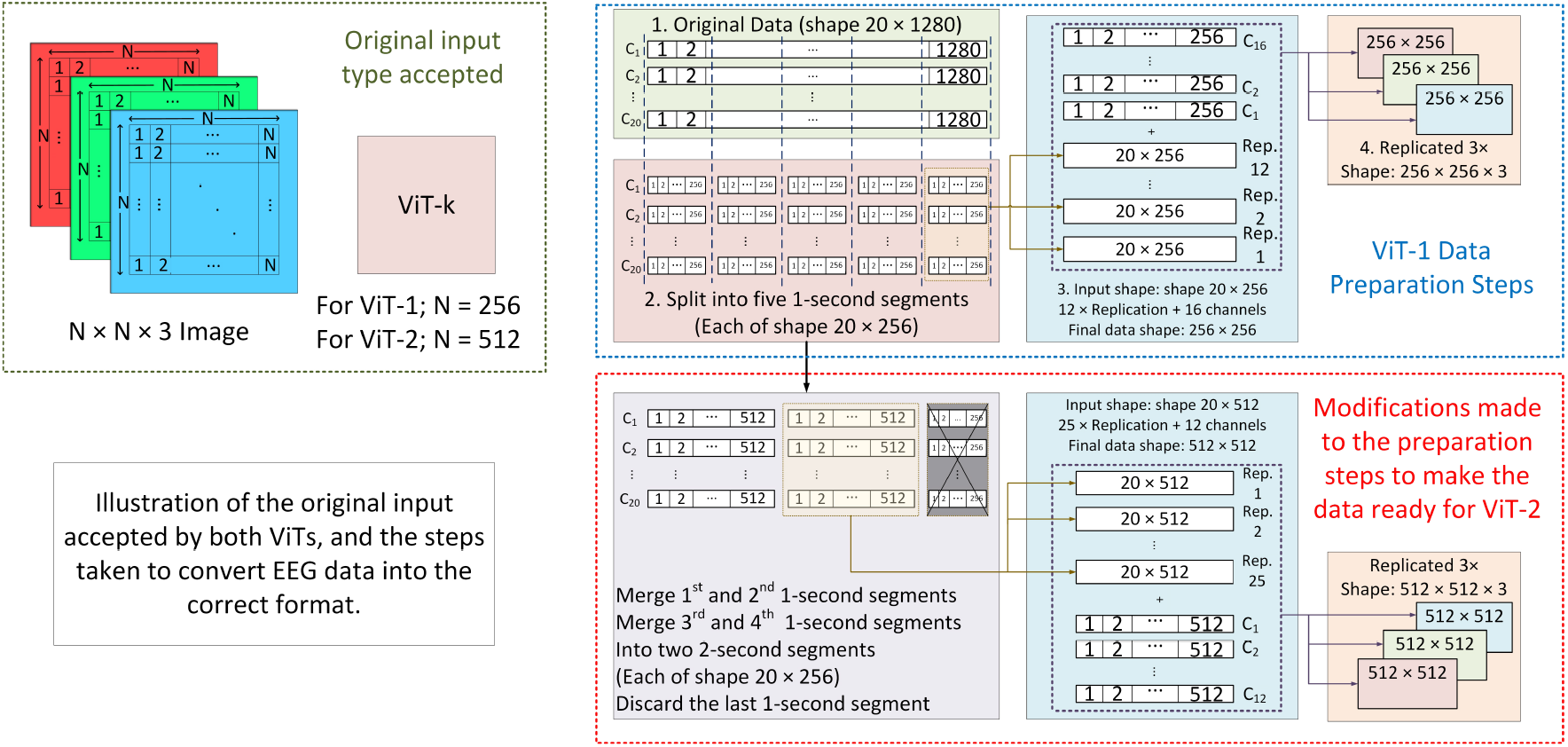
Illustration of the phases in which data is augmented to make it suitable for being input into a ViT.

### A. ViT-1 Design

ViT-1 consists of six stages: stage one performs patch partitioning, the next four stages are transformer blocks followed by the sixth stage which is the classifier head. While most transformers use a two-step strategy to calculate both local and global self-attention, we have used a modified version of [13] which has these key differences [14]: the attention blocks apply a scaled cosine function to the query and key, log-spaced coordinates are used in the dense layers, and the normalization layer is moved to the end of each residual unit. The input stage of ViT-1 stage accepts an RGB image of shape *H* ×*W* ×3, though the recommended shapes are either *N* ×*N* ×3 with *N* = 256 or *N* = 512. The first stage partitions each image to *H/*4×*W/*4 patches and the embedding layer projects it onto a feature dimension *C* (*C* is a hyperparameter). From stages two to five, the patch size progresses as follows: *H/*4×*W/*4×*C* → *H/*8×*W/*8×2*C* → *H/*16×*W/*16×4*C* → *H/*32×*W/*32×8*C*. The classifier head performs 1D average pooling on the *H/*32 ×*W/*32 tokens followed by dense layers.

The major innovations we performed include formatting the data so that it is suitable for the patch partitioning phase and the classifier head to make it suitable for binary classification. Recall that the multi-channel EEG time-series data samples are available in the shape 20 × 1280 whereas the pre-trained model checkpoint used accepts 256 × 256 × 3. Therefore, we first reduce the window size from five seconds to one second, which splits each segment further into five data points of shape 20 × 256; the five new data points share the same label (preictal or interictal) as the original data of shape 20 × 1280. Hence, the dataset size of each benchmark grows five-fold albeit each preictal or interictal sample has a smaller resolution of one second. To increase the size along the channel dimension, we over-sample the 20 channels by replicating them 12 times resulting in data of shape 240 × 256, and then 16 channels are over-sampled randomly to create a 256 × 256 image-like representation. We feed the same image to each of the RGB channels of the model. The classifier head is re-designed to predict two classes and added with randomly initialized weights. As part of our partial re-training strategy, we only train stage 1 (up-to the embedding layer), the normalization layer before the classifier and the classifier itself with the following hyper parameters: batch size of 128, 5 epochs, learning rate (LR) of 1*e* − 3, a warm-up ratio of 0.1, and a 1% weight decay. The warm-up ratio is the ratio of the required training steps needed to reach the pre-set learning rate of 1*e* − 3 to the total steps in 1 epoch. From stage 2 transformer layer up-to stage 5 are frozen.

### B. ViT-2 Design

ViT-2 is adopted from the Segformer [15] and also has six stages similar to ViT-1. Each transformer block in stages 2-5 contains a patch merging layer, a transformer comprising both an encoder and a decoder, and the normalization layer as illustrated in Fig. 2. Instead of positional embeddings, the encoder uses a combination of dense layers and 3 × 3 convolutions to get positional information *Y* in two steps.

**Fig. 2.**
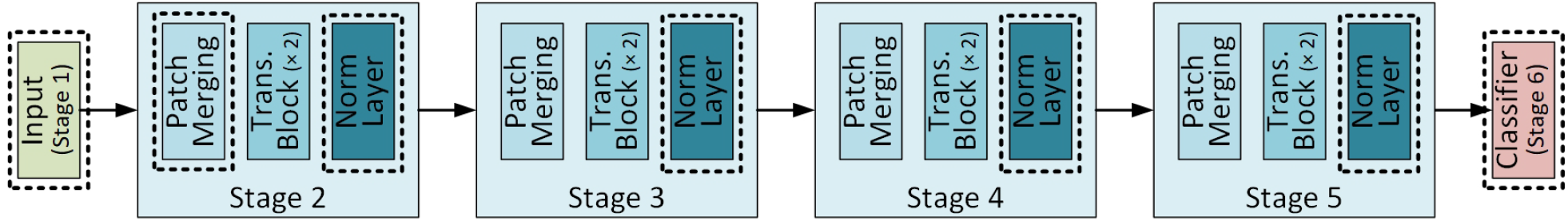
Illustration of the different stages of ViT-2 with the re-trained layers encapsulated by dotted rectangles.

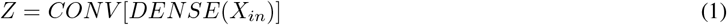

where *X*_*in*_ is the feature from self-attention modules, *DENSE* represents the fully connected (FC) network, and *CONV* is the convolution operation with a 3 *×* 3 kernel.

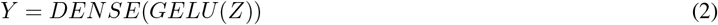

where *GELU* is the Gaussian Error Linear Unit activation function. The first transformer block in stage 2 uses a 2D convolution kernel 7×7, stride 4 and padding 3×3 for patch merging. In stages 3-5, a kernel of size 3×3 with stride 2 and padding 1×1 is used. The pretrained model used is from the NVIDIA/Mit-B0 checkpoint, which is only based on the encoder, is fine-tuned on ImageNet-1k and works with input images of size 512 × 512 × 3.

To make the EEG data ready for ViT-2, we reduce the window size from five to two seconds and discard one second of data. Therefore, the original 5-second segment is split into two chunks each of shape 20×512; each shares the same label as the original data of shape 20 × 1280. Finally, each data sample is replicated 25 times along the y-axis to get data of shape 500 × 512. An additional 12 randomly sampled channels are added and the data is replicated 3 times to create the final data of shape 512×512×3. The classifier head is re-designed to predict two classes. As part of the re-training strategy, we choose to re-train the patch merging layer of stage 2, the normalization layers of each block and the classifier layer (as highlighted in Fig. 2) with the following hyperparameters: batch size of 128, 5 epochs, LR set to 1*e* − 3, a warm-up ratio of 0.1 and 1% weight decay.

### C. LLM (Longformer)

To test the usability of an LLM for seizure prediction, we used a pre-trained version of the LongFormer [12] which can process up-to 4,096 tokens. The main underlying mechanism involves two types of “attention patterns”. First, a sliding window of size 512 is used to compute the attention of each token, and multiple window-based attention scores are stacked to form the top layer which cycles over all input tokens. Symmetric global attention is then used to for each pair of tokens in a given sequence followed by classification using a ‘CLS’ token. The major modification we performed on the data is converting a sequence of real numbers to a sequence of string tokens. An example token from EEG data appears as follows: {‘-’, ‘3’, ‘.’, ‘714’, ‘307’, ‘22’, ‘37’, ‘31’, ‘193’, ‘e’, ‘-’, ‘05’, ‘Ġ -’, ‘2’, ‘.’, ‘07’, ‘465’, ‘80’, ‘889’, ‘309’, ‘362’, ‘e’, ‘-’, ‘05’, ‘Ġ 1’, ‘.’, ‘606’, ‘18’, ‘690’, ‘75’, ‘594’, ‘348’}.

Tokenizing the original EEG segment of shape 20 × 1280 results in more than 25,000 tokens whereas only a maximum of 4,096 are allowed by the LLM. Reducing the window size to 1 second (256 samples) also does not help. Hence, we apply an innovative new approach inspired by [4] where each of the 20 channels are tokenized individually. First, we convert each data point of shape 20 × 256 into 20 sequences of length 256, each with the same label as the original data, and extract tokens from each sequence and set an upper limit of 2,500 tokens. The rationale behind string tokenization is to make the EEG data LLM-ready, the rationale for using single-channel sequences include better performance and the ability to train without the exhaustion of resources and the rationale for shorter segments is to stay within memory bounds and simultaneously avoid over-fitting. We train only the classifier layer using the following hyperparameters: LR of 5*e* − 4, warm-up ratio 0.03, weight decay of 1%, batch size 64 and train for 4 epochs. The model makes one prediction for each channel but the final decision is based on majority voting, where a seizure (preictal label) is predicted only if more than 10 channels indicate a positive label.

### D. Data and Evaluation

We used our recently introduced patient-independent seizure prediction benchmark dataset *MLSPred-Bench* [5]. Recall that seizure prediction requires identifying preictal segments. Therefore, the benchmark dataset extracts preictal segments from a pre-defined seizure prediction horizon (SPH) after allowing for a gap time equal to the seizure occur(rence) period (SOP) before the start of a seizure. However, because preictal phases are not clinically identifiable on an EEG, we test a combination of values for the SPH and SOP based on the literature, where *SPH* ∈ {2, 5, 15, 30} minutes and the *SOP* ∈ {1, 2, 5} minutes. As observable, this leads to 12 benchmarks as illustrated in Table I. More information on the benchmarks can be read in our recent paper [5].

**TABLE I.** Illustration of the benchmarks

| SPH/SOP | 1 | 2 | 5 |
| --- | --- | --- | --- |
| 2 | BM1 | BM2 | BM3 |
| 5 | BM4 | BM5 | BM6 |
| 15 | BM7 | BM8 | BM9 |
| 30 | BM10 | BM11 | BM12 |

For evaluation, let us define the true positive (*TP*) as a correct seizure prediction and a false positive (*FP*) as an incorrect seizure prediction. Correctly predicting no seizure is a true negative (*TN*) whereas incorrectly predicting no seizure is a false negative (*FN*). Then, the ability of the model to correctly predict a seizure is measured by the sensitivity (*Sen*):

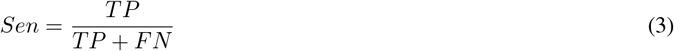

The ability of an ML model to correctly identify the negative class is given by the specificity (*Spe*):

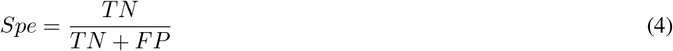

The overall ability of the model to predict the correct labels is given by the accuracy (*Acc*):

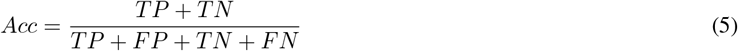

Along with these 3 metrics, we can evaluate the receiver operating characteristic (ROC) - area under the curve (AUC) scores which measures the ability of the model to offer a trade-off between the sensitivity and specificity. In the next section, we use these metrics to compare both ViTs and the LLMs for all benchmarks against baseline results from a residual neural network (ResNet)-based model [18] available in [5].

## IV. Results and Discussions

Table II compares the AUC scores for both ViTs and the LLM against SPERTL [18]. From a cursory glance, we can interpret that ViT-2 outperforms all other models for the first six benchmarks which use a maximum SPH (preictal time) of five minutes. The highest ROC is 77.00% on benchmark (BM) 6 - (BM6), whereas ResNet has the highest AUC scores from BM7 to BM9 which use a SPH of 15 minutes. However, Longformer was able to beat every other model from BM10 to BM12 with the highest AUC score of 87.57% on BM11. This is also the highest we have observed across all our experiments. On average, the LLM does provide the overall highest average AUC score which we focus on, because it quantifies the ability of the model to provide a high accuracy while providing a good trade-off between the sensitivity and specificity.

**TABLE II.** Comparison of AUC-ROC Scores for all models validated on all benchmarks.

| BM | ViT-1 | ViT-2 | LLM | ResNet |
| --- | --- | --- | --- | --- |
| 1 | 69.86% | **76.30%** | 69.45% | 66.80% |
| 2 | 72.64% | **76.20%** | 69.79% | 66.50% |
| 3 | 73.32% | **76.10%** | 73.22% | 69.40% |
| 4 | 73.13% | **76.70%** | 70.23% | 63.00% |
| 5 | 71.81% | **76.06%** | 71.58% | 68.70% |
| 6 | 73.65% | **77.00%** | 71.08% | 72.90% |
| 7 | 73.04% | 71.91% | 69.30% | **81.10%** |
| 8 | 71.43% | 71.95% | 70.81% | **83.60%** |
| 9 | 74.18% | 71.23% | 71.83% | **79.20%** |
| 10 | 43.50% | 41.00% | **62.07%** | 56.40% |
| 11 | 80.50% | 79.22% | **87.57%** | 80.50% |
| 12 | 45.03% | 42.77% | **60.41%** | 50.80% |
| Avg. | 68.51% | 69.70% | **71.40%** | 70.91% |

However, an AUC score does not necessarily guarantee a high prediction sensitivity or specificity. Therefore, we compare the sensitivity, accuracy and specificity of each model for all benchmarks in Table III. The first interesting result is that the AUC scores on their own do not necessarily translate to high predictive power when using a classification threshold of 0.5. For example, even though the ROC-AUC score of ViT-2 was better than every other model from BM1 to BM6, the LLM actually achieves a higher accuracy and sensitivity compared to all other models on all BMs except for BM2 and BM6. For example, ViT-2 has a 70.74% accuracy for BM2 whereas the LLM has an accuracy of 70.34%. Similarly, ViT-2 has a 76.10% sensitivity on BM6 whereas the LLM has a sensitivity of 66.98%. Most of results show higher sensitivity than specificity, which implies that using only EEG data, the ViTs and LLM used lean towards predicting a seizure with a threshold of 0.5 despite the problem being made class-balanced. Though the specificity can be improved by modifying the threshold, it will reduce the sensitivity which is more significant as missed seizures are more critical.

**TABLE III.** Accuracy, specificity and sensitivity of each model across all benchmarks.

| BM | ViT-1 |  |  | ViT-2 |  |  | LLM |  |  | ResNet |  |  |
| --- | --- | --- | --- | --- | --- | --- | --- | --- | --- | --- | --- | --- |
|  | Acc | Sen | Spe | Acc | Sen | Spe | Acc | Sen | Spe | Acc | Sen | Spe |
| 1 | 66.41% | 75.50% | 57.32% | 69.70% | 66.50% | 72.80% | 71.48% | 79.95% | 63.18% | 61.83% | 90.81% | 32.85% |
| 2 | 68.16% | 66.76% | 69.56% | 70.74% | 73.90% | 67.50% | 70.34% | 82.83% | 57.57% | 50.02% | 99.67% | 0.37% |
| 3 | 68.47% | 76.26% | 60.68% | 68.60% | 69.57% | 67.79% | 74.30% | 85.48% | 63.14% | 55.05% | 98.37% | 11.72% |
| 4 | 65.65% | 71.68% | 59.63% | 67.50% | 73.50% | 61.50% | 71.16% | 82.38% | 59.81% | 57.79% | 63.18% | 52.40% |
| 5 | 67.21% | 73.00% | 61.42% | 68.27% | 73.79% | 62.74% | 70.86% | 78.94% | 62.46% | 57.39% | 29.06% | 85.72% |
| 6 | 67.74% | 75.03% | 60.46% | 70.30% | 76.10% | 64.50% | 71.26% | 66.98% | 75.54% | 60.02% | 22.94% | 97.09% |
| 7 | 67.30% | 76.19% | 58.42% | 66.98% | 75.18% | 58.78% | 67.60% | 84.34% | 50.65% | 52.98% | 6.97% | 99.00% |
| 8 | 65.82% | 76.19% | 55.44% | 65.25% | 71.00% | 59.44% | 70.52% | 86.54% | 54.44% | 50.99% | 2.64% | 99.33% |
| 9 | 67.22% | 75.33% | 59.11% | 64.68% | 69.42% | 59.90% | 73.48% | 89.91% | 57.54% | 49.90% | 99.58% | 0.23% |
| 10 | 48.43% | 61.60% | 35.27% | 46.39% | 66.77% | 26.00% | 66.60% | 66.99% | 66.24% | 50.00% | 100.00% | 0.00% |
| 11 | 70.04% | 63.93% | 76.14% | 71.18% | 64.87% | 77.48% | 77.87% | 66.73% | 89.01% | 50.00% | 100.00% | 0.00% |
| 12 | 48.90% | 61.10% | 36.70% | 50.13% | 63.19% | 37.08% | 64.10% | 77.17% | 50.84% | 50.00% | 100.00% | 0.00% |
| Avg | 64.28% | 71.05% | 57.51% | 64.98% | 70.32% | 59.63% | **70.80%** | **79.02%** | **62.53%** | 53.83% | 67.77% | 39.89% |

### A. Discussion on why some models work better

The ability of the ViT to provide a good AUC score for the first six benchmarks shows that it can make accurate predictions based on small preictal times. One characteristic of the first six benchmarks is that short-term local data was used as features. Whereas the window duration was 5 seconds across all 12 benchmarks implying 1280 temporal features, the number of preictal and interictal windows for each seizure were considerably smaller per seizure compared to benchmarks 7-12. Hence, the ViTs work better because the underlying architecture is efficient at extracting the spatial features along fewer short-term local temporal features, but are unable to discern long-term temporal dependencies or correlations when a longer preictal duration are selected. Once we move to benchmarks 7-9, the number of preictal samples per seizure are increased by 200%, but the dataset size is smaller (with fewer seizures). This data may be too small for much deeper architectures such as ViTs and LLMs but ideal for ResNets which are ‘less’ deep [19]. Hence, ResNet provides a higher AUC score for benchmarks 7-9. The best AUC score for benchmarks 10-12 which have the highest SPH is provided by the LLM followed by ResNet and then ViT. This is due to the combined impact of tokenizing the sequences per channel and doubling of the number of preictal and interictal samples per seizure which increases the dataset size sufficiently for LLMs.

### B. Experiments for Ablation Study, Generalization and ML

Recall that for both ViT-1 and ViT-2, we unfroze the patch embedding in stage 1, the normalization layer before classification and classifier head in stage 6. For the ablation studies, we tried re-training and testing the models with different combinations of layers frozen. Table IV shows the ROC-AUC scores for benchmarks 5 and 8. It is observable that re-training all three stages clearly provides a significantly higher AUC score compared to when only one or two stages are re-trained.

**TABLE IV.** Ablation Study: ROC-AUC Scores for VIT-1 and VIT-2 with different configurations of re-trained layers.

| Model | ViT-1 |  | ViT-2 |  |
| --- | --- | --- | --- | --- |
| Re-trained Layers | BM5 | BM8 | BM5 | BM8 |
| Classifier Only | 61.21% | 59.12% | 69.37% | 65.67% |
| Classifier+Norm | 61.67% | 58.50% | 71.35% | 68.73% |
| Patch Embedding | 70.38% | 67.39% | 68.51% | 70.82% |
| All three | **71.81%** | **71.43%** | **76.06%** | **71.95%** |

Table V shows the superiority of the proposed techniques which improve the AUC score and accuracy by at least 4% compared to four traditional ML methods including: logistic regression (LR), k-Nearest Neighbor (kNN), adaptive boosting (AdaBoost) and bagging. To show the ability of the model to generalize, we test it on another EEG dataset from subjects with Alzheimer’s Disease (AD) where the task was to discriminate between segments from healthy controls and subjects with AD. As the results indicate, both ViT-1 and ViT-2 worked well with ViT-2 outperforming ViT-1 by about 8% in terms of AUC score, accuracy by 7%, sensitivity by 8% and specificity by more than 6%.

**TABLE V.**
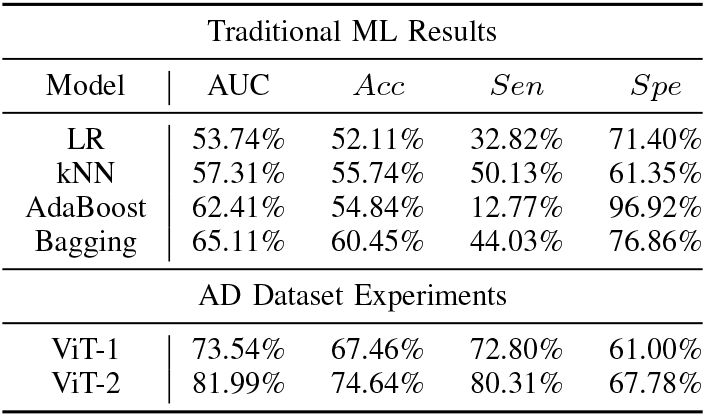
Results for Traditional ML and Generalization on AD

### C. Computational Resource Discussion

Our experiments were carried out on our in-house high performance computing (HPC) clusters Raptor and Dragon. Raptor has two HPC central processing units (CPUs) and is equipped with one Graphical Processing Unit (GPU) with 3,849 cores and 12GB memory. Dragon has 6 compute nodes and each node has two 10-core CPUs and 2 GPUs with 10,752 cores each and up-to 96GB total memory. All data wrangling was done on the Dragon compute nodes, the ML experiments on the Raptor compute nodes, the DL experiments with ResNet on Raptor GPU and the VIT and LLM experiments each were done on 1 Dragon node, i.e., on 2 GPUs each. Despite the higher resources used by the ViT and LLMs, on average the ViTs required nearly half an hour of training time per benchmark for 5 epochs whereas the LLM required up-to 60 hours for 4 epochs. In contrast, the ResNet required around 12 minutes per benchmark for 25 epochs whereas the ML models only required a few seconds with the exception of AdaBoost which required 5 minutes.

## V. Conclusion and Future Work

In this work, we developed strategies to reformulate imaging and textual data pre-trained models (2 vision transformers and one LLM) into methods that can process multidimensional time-series EEG data. Our evaluation was accomplished using recent benchmarking datasets, called MLSPred-Bench [5]. Our results indicate that the LLM has the best performance for all metrics by achieving a sensitivity of 79.02%, a specificity of 62.53%, and an accuracy of 70.80%, highlighting its potential as a robust tool for seizure prediction in clinical and home-care settings. Future work includes stratification of different model stages for performance, and data augmentation techniques for better sensitivity. Novel transformer models are also part of the future investigation.

## References

[1] World Health Organization, “Epilepsy,” 2023.

[2] N. Mühlenfeld, P. Stürmann, I. Marzi, F. Rosenow, A. Strzelczyk, R. D. Verboket, and L. M. Willems, “Seizure related injuries – Frequent injury patterns, hospitalization and therapeutic aspects,” Chinese Journal of Traumatology - English Edition, vol. 25, pp. 272–276, sep 2022.

[3] A. H. Shoeb and J. Guttag, “Application of Machine Learning To Epileptic Seizure Detection,” in ICML, pp. 975–982, jan 2010.

[4] Z. Wang, J. Yang, H. Wu, J. Zhu, and M. Sawan, “Power efficient refined seizure prediction algorithm based on an enhanced benchmarking,” Scientific Reports, vol. 11, ec 2021.

[5] U. Mohammad and F. Saeed, “MLSPred-Bench: ML-Ready Benchmark Leveraging Seizure Detection EEG data for Predictive Models,” bioRxiv preprint bioRxiv:2024.07.17.604006, pp. 1–13, 2024.

[6] H. Daoud and M. A. Bayoumi, “Efficient Epileptic Seizure Prediction Based on Deep Learning,” IEEE Transactions on Biomedical Circuits and Systems, vol. 13, pp. 804–813, oct 2019.

[7] Y. Ma, Z. Huang, J. Su, H. Shi, D. Wang, S. Jia, and W. Li, “A Multi-Channel Feature Fusion CNN-Bi-LSTM Epilepsy EEG Classification and Prediction Model Based on Attention Mechanism,” IEEE Access, vol. 11, pp. 62855–62864, 2023.

[8] X. Lu, A. Wen, L. Sun, H. Wang, Y. Guo, and Y. Ren, “An Epileptic Seizure Prediction Method Based on CBAM-3D CNN-LSTM Model,” IEEE Journal of Translational Engineering in Health and Medicine, vol. 11, pp. 417–423, 2023.

[9] L. Guo, T. Yu, S. Zhao, X. Li, X. Liao, and Y. Li, “CLEP: Contrastive Learning for Epileptic Seizure Prediction Using a Spatio-Temporal-Spectral Network,” IEEE Transactions on Neural Systems and Rehabilitation Engineering, vol. 31, pp. 3915–3926, 2023.

[10] L. Xia, R. Wang, H. Ye, B. Jiang, G. Li, C. Ma, and Z. Gao, “Hybrid LSTM-Transformer Model for the Prediction of Epileptic Seizure Using Scalp EEG,” IEEE Sensors Journal, vol. 24, pp. 21123–21131, jul 2024.

[11] S. Shi and W. Liu, “B2-ViT Net: Broad Vision Transformer Network with Broad Attention for Seizure Prediction,” IEEE Transactions on Neural Systems and Rehabilitation Engineering, vol. 32, pp. 178–188, 2024.

[12] I. Beltagy, M. E. Peters, and A. Cohan, “Longformer: The Long-Document Transformer,” arXiv preprint arXiv:2004.05150, pp. 1–17, apr 2020.

[13] A. Dosovitskiy, L. Beyer, A. Kolesnikov, D. Weissenborn, X. Zhai, T. Unterthiner, M. Dehghani, M. Minderer, G. Heigold, S. Gelly, J. Uszkoreit, and N. Houlsby, “An Image is Worth 16×16 Words: Transformers for Image Recognition at Scale,” in International Conference on Learning Representations, pp. 1–21, 2021.

[14] Z. Liu, H. Hu, Y. Lin, Z. Yao, Z. Xie, Y. Wei, J. Ning, Y. Cao, Z. Zhang, L. Dong, F. Wei, and B. Guo, “Swin Transformer V2: Scaling Up Capacity and Resolution,” Proceedings of the 2022 IEEE/CVF Conference on Computer Vision and Pattern Recognition (CVPR), pp. 11999–12009, 2022.

[15] E. Xie, W. Wang, Z. Yu, A. Anandkumar, J. M. Alvarez, and P. Luo, “SegFormer: Simple and Efficient Design for Semantic Segmentation with Transformers,” in 2021 Conference on Neural Information Processing Systems (NeurIPS 2021) (A. Beygelzimer, Y. Dauphin, P. Liang, and J. W. Vaughan, eds.), pp. 1–14, 2021.

[16] T. Dissanayake, T. Fernando, S. Denman, S. Sridharan, and C. Fookes, “Deep Learning for Patient-Independent Epileptic Seizure Prediction Using Scalp EEG Signals,” IEEE Sensors Journal, vol. 21, pp. 9377–9388, apr 2021.

[17] T. Dissanayake, T. Fernando, S. Denman, S. Sridharan, and C. Fookes, “Geometric Deep Learning for Subject-Independent Epileptic Seizure Prediction using Scalp EEG Signals,” IEEE Journal of Biomedical and Health Informatics, vol. 26, no. 2, pp. 527–538, 2021.

[18] U. Mohammad and F. Saeed, “SPERTL: Epileptic Seizure Prediction using EEG with ResNets and Transfer Learning,” in BHI-BSN 2022 - IEEE-EMBS International Conference on Biomedical and Health Informatics and IEEE-EMBS International Conference on Wearable and Implantable Body Sensor Networks, Symposium Proceedings, pp. 1–5, Institute of Electrical and Electronics Engineers Inc., 2022.

[19] H. Ismail Fawaz, G. Forestier, J. Weber, L. Idoumghar, and P. A. Muller, “Deep learning for time series classification: a review,” Data Mining and Knowledge Discovery, vol. 33, pp. 917–963, jul 2019.

